# Incomplete MyoD-induced transdifferentiation is associated with chromatin remodeling deficiencies

**DOI:** 10.1101/171470

**Authors:** Dinesh Manandhar, Lingyun Song, Ami Kabadi, Jennifer B. Kwon, Lee E. Edsall, Melanie Ehrlich, Koji Tsumagari, Charles Gersbach, Gregory E. Crawford, Raluca Gordan

## Abstract

Our current understanding of cellular transdifferentiation systems is limited. It is oftentimes unknown, at a genome-wide scale, how much transdifferentiated cells differ quantitatively from both the starting cells and the target cells. Focusing on transdifferentiation of primary human skin fibroblasts by forced expression of myogenic transcription factor MyoD, we performed quantitative analyses of gene expression and chromatin accessibility profiles of transdifferentiated cells compared to fibroblasts and myoblasts. In this system, we find that while many of the early muscle marker genes are reprogrammed, global gene expression and accessibility changes are still incomplete when compared to myoblasts. In addition, we find evidence of epigenetic memory in the transdifferentiated cells, with reminiscent features of fibroblasts being visible both in chromatin accessibility and gene expression. Quantitative analyses revealed a continuum of changes in chromatin accessibility induced by MyoD, and a strong correlation between chromatin-remodeling deficiencies and incomplete gene expression reprogramming. Classification analyses identified genetic and epigenetic features that distinguish reprogrammed from non-reprogrammed sites, and suggested ways to potentially improve transdifferentiation efficiency. Our approach for combining gene expression, DNA accessibility, and protein-DNA binding data to quantify and characterize the efficiency of cellular transdifferentiation on a genome-wide scale can be applied to any transdifferentiation system.

## INTRODUCTION

MyoD is a master transcription factor (TF) critical for normal muscle development. Overexpression of this single TF has been shown to transdifferentiate cells from non-muscle lineages, such as dermal fibroblasts, chondrocytes and adipocytes, into cells with muscle-like expression and phenotypic characteristics (1-4). Expression studies on MyoD-transdifferentiated cells show that MyoD overexpression upregulates many myogenic genes, including endogenous MyoD by an autoregulatory loop (1,2,5). However, some studies suggest that not all genes involved in muscle-specific functions are upregulated in response to MyoD overexpression (6,7). In addition, as with other transdifferentiation studies, genes and pathways specific to the donor cells do not get completely suppressed during TF-induced transdifferentiation (2,8). In this study, we use MyoD-induced transdifferentiation of fibroblasts as a model system to develop a general approach to characterize and study transdifferentiation deficiencies at the chromatin and gene expression level, with the goal of generating testable hypotheses for how the transdifferentiation process could be improved.

A major limitation of previous MyoD-induced transdifferentiation studies is that only a handful of myogenic markers were analyzed (1,2,9,10). Microarray experiments have also been used to analyze gene expression changes during MyoD-induced reprogramming (6,11), but even these studies were limited to a few thousand genes. Recently, it has been recognized that more comprehensive genome-wide analyses (e.g., RNA-seq and ChIP-seq) are critical to assess whether full or partial reprogramming is achieved by overexpression of MyoD or other myogenic regulatory factors (MRFs) (12). A few recent studies analyzed genome-wide changes induced by MyoD overexpression, but they focused either on comparing gene expression changes induced by wild-type versus engineered MyoD variants (13), on MyoD binding events in transdifferentiated cells versus myoblasts and myotubes in mouse (14), or on comparing the sets of genes upregulated during myogenic versus neurogenic transdifferentiation of mouse fibroblast cells (15). To our knowledge, a direct genome-wide comparison of the gene expression profiles of MyoD-transdifferentiated cells against primary human muscle cells has not been performed.

In addition, as with many other transdifferentiation systems, previous MyoD-induced reprogramming studies also have not characterized chromatin accessibility changes to better understand the transdifferentiation process (16,17). This is particularly relevant for MyoD-induced cellular reprogramming because (i) MyoD is a known pioneer factor capable of remodeling chromatin and changing the regulatory landscape (18), and (ii) it has been suggested that the limited ability of MyoD to reprogram certain cell types, such as P19 mouse embryonal carcinoma cell line, human HepG2, HeLa, and undifferentiated ES cells, is likely attributable to starting cell type-specific differences in chromatin landscape (12,19).

We have generated and analyzed global gene expression (RNA-seq), chromatin accessibility (DNase-seq), and MyoD binding (ChIP-seq) data on human primary skin fibroblasts transduced with an inducible FLAG-tagged human MyoD expression cassette. To assess the efficiency of the transdifferentiation process, we compared these data to gene expression, chromatin accessibility, and MyoD binding profiles generated from primary human myoblasts and primary skin fibroblasts. Our study reveals that MyoD overexpression leads to a continuum of DNaseI hypersensitive (DHS) site changes. Some DHS sites were completely reprogrammed, i.e the chromatin accessibility at those sites in transdifferentiated cells closely resembles the accessibility in primary myoblasts. However, many other sites are either not reprogrammed or partially reprogrammed. Using a classification approach to analyze reprogrammed and non-reprogrammed DHS regions, we identify potential explanations for the incomplete reprogramming at the chromatin level and we suggest ways to improve the process through additional regulatory factors and induced epigenetic changes. Finally, we provide evidence of a strong correlation between chromatin reprogramming deficiencies and lack of complete reprogramming of gene expression. Given the causal role of chromatin accessibility in regulating gene expression, our analyses suggest that incomplete MyoD transdifferentiation is due, at least in part, to deficiencies in remodeling the chromatin landscape.

## METHODS

### Cell culture

Primary human dermal fibroblasts (Catalog ID: GM03348) were obtained from Coriell Institute (Camden, New Jersey) and were maintained in DMEM supplemented with 10% FBS and 1% penicillin streptomycin. All cells were cultured at 37 °C with 5% CO2. Three biological replicates of both the parental fibroblast cells and transduced fibroblast cells were harvested on different days. Four replicates of myoblasts were originally sampled from biopsies on quadriceps and grown as described and published previously (20-22). As described previously, myoblasts cells were harvested at ~70% confluence, and all myoblast preparations had <1% multinucleated cells. Importantly, immunostaining of the myoblast samples showed 90-98% desmin-positive cells (21), indicative of highly pure populations of myoblasts.

### Viral production and transduction

All lentiviral vectors used in this study were produced from second-generation plasmids using standard viral production methods previously described (13). Briefly, 3.5 million HEK293T cells were plated per 10cm dish. The following day, cells were transfected with 20 μg of transfer vector, 6 μg of pMD2G, and 10 μg psPAX2 using a calcium phosphate transfection. The media was changed 12-14 hours post transfection. The viral supernatant was collected 24 and 48 hours after this media change and pooled. For transduction, the cell medium was replaced with viral supernatant supplemented with 4 μg/mL polybrene. The viral supernatant was changed 24-48 hours later.

### MyoD-directed genetic reprogramming

Human dermal fibroblasts were transduced with a Tet-ON lentivirus that expresses a 3xFlag-tagged full-length *MYOD1* cDNA (13). In this vector, a 3xFlag-tagged full-length human *MYOD1* cDNA-T2A-dsRed-Express2 cassette is expressed from the Tetracycline Responsive Element (TRE) promoter. T2A is a peptide that facilitates ribosomal skipping as the mRNA transcript is being translated into protein. The resulting products are two separate peptides that are expressed in a similar ratio. The vector constitutively expresses the Reverse Tetracycline Transactivator (rtTA2s-M2) and the Puromycin resistance gene from the human phosphoglycerate kinase (hPGK) promoter. The rtTA and Puro^R^ are co-expressed from the same mRNA via an Internal Ribosomal Entry Site (IRES). The rtTA is only able to bind to the TRE and activate expression of the downstream genes in the presence of doxycycline.

Transduced cells were selected in 1 μg/ml puromycin for 4 days until 100% cell death was observed in untransduced cells, in order to obtain a pure population. Cells were expanded in standard growth medium. Selected cells were seeded in appropriate plates and grown to confluence. 3xFlag-tagged MyoD transgene expression was induced by supplementing the medium with 3 μg/mL doxycycline. Cells were given fresh media supplemented with doxycycline every two days. All differentiation studies were conducted in standard growth medium (DMEM supplemented with 10% FBS and 1% penicillin streptomycin). Cells were harvested after 10 days of doxycycline treatment. These cells are referred to subsequently as MyoD-induced. Given that cells were not transferred to differentiation media to further induced terminal differentiation, in all our computational analyses we compare the transdifferentiated cells against myoblasts, which represent the early myogenic determination stage.

### Quantitative reverse transcription PCR

RNA was isolated using the RNeasy Plus RNA isolation kit (Qiagen). cDNA synthesis was performed using the SuperScript VILO cDNA Synthesis Kit (Invitrogen). Real-time PCR using PerfeCTa SYBR Green FastMix (Quanta Biosciences) was performed with the CFX96 Real-Time PCR Detection System (Bio-Rad). Oligonucleotide primers and PCR conditions are reported in **Table S1**. Primer specificity was confirmed by agarose gel electrophoresis and melting curve analysis. Reaction efficiencies over the appropriate dynamic range were calculated to ensure linearity of the standard curve (13). The results are expressed as fold-increase mRNA expression of the gene of interest normalized to *Beta Actin* expression using the ΔΔCt method. Reported values are the mean and S.E.M. from two independent experiments (n = 2) where technical replicates were averaged for each experiment. Effects were evaluated with multivariate ANOVA and Dunnett’s post hoc test using JMP 10 Pro.

### Western blot

Cells were lysed in RIPA Buffer (Sigma) supplemented with protease inhibitor cocktail (Sigma). Protein concentration was measured using BCA protein assay reagent (Thermo Scientific) and BioTek Synergy 2 Multi-Mode Microplate Reader. Lysates were mixed with loading buffer and incubated at 70°C for 5 min; equal amounts of protein were run in NuPage 10% Bis-Tris Gel polyacrylamide gels (Bio-Rad) and transferred to nitrocellulose membranes. Nonspecific antibody binding was blocked with TBST (50 mM Tris, 150 mM NaCl and 0.1% Tween-20) with 5% nonfat milk for 1hour at room temperature. The membranes were incubated with primary antibodies (anti-MyoD (Santa Cruz, Sc-32758) in 5% BSA in TBST, diluted 1:250, overnight at 4°C; anti-Myogenin (Santa Cruz, Sc-12732) in 5% BSA, diluted 1:250, overnight at 4°C; anti-Beta Actin (Sigma, A2066) in 5% milk in TBST, diluted 1:5,000, for 30 min at room temperature, and the membranes were washed with TBST for 15 min. Membranes were incubated with anti-rabbit HRP-conjugated antibody (Sigma, A 6154) or anti-mouse HRP-conjugated antibody (Santa Cruz, SC-2005) diluted 1:5,000 for 30 min and washed with TBST for 15 min. Membranes were visualized using the ImmunStar WesternC Chemiluminescence Kit (Bio-Rad) and images were captured using a ChemiDoc XRS+ System and processed using ImageLab software (Bio-Rad).

### Immunofluorescence staining

Fibroblasts transduced with Tet-ON LV co-expressing 3xFlag human MyoD and dsRed Express2 were plated on autoclaved glass coverslips (1mm, Thermo Scientific). Following 10 days of transgene expression, cells were fixed in 4% PFA and prepared for immunofluorescence staining. Samples were permeabilized in blocking buffer (PBS supplemented with 5% BSA and 0.2% Triton X-100) for one hour at room temperature. Samples were incubated with primary antibodies MF20 (Hybridoma Bank) diluted 1:200 and MyoD (Santa Cruz, Sc-32758) diluted 1:250 in blocking buffer overnight at 4°C, and rinsed for 15 min in PBS. Samples were incubated with secondary antibodies anti-mouse IgG2b Alexa Fluor 488 (Invitrogen, A-21141) and anti-mouse IgG1 Alex Fluor 647 (Invitrogen, A-21240) both diluted 1:500 for one hour at room temperature. Cells were incubated with DAPI diluted 1:5000 in PBS for 5 min and washed with PBS for 15 min. Coverslips were mounted with ProLong Gold Antifade Reagent (Invitrogen) and imaged using a Leica SP5 inverted confocal microscope.

### Flow cytometry analysis

Following 8 days of transgene expression, cells were analyzed for dsRed expression by flow cytometry. Untransduced fibroblasts and fibroblasts transduced with Tet-ON LV co-expressing 3xFlag human MyoD and dsRed Express2 were harvested, washed once with PBS, and resuspended in 3% FBS in PBS. All cells were analyzed using the SH800 Cell Sorter (Sony Biotechnology).

### DNase-seq and defining fibroblast- or myoblast-specific DHS sites

DNase-seq was performed as previously described (23), with one modification: oligo 1b was synthesized with a 5’ phosphate to increase the efficiency of ligation. 5-20 million cells were used for each biological replicate. Libraries were sequenced on Illumina GAII or Hi-Seq 2000 sequencing platform in Duke Sequencing and Analysis Core Resource. Raw reads were trimmed to 20bp from 5’ and aligned to hg19 reference genome by using bowtie-0.12.9, with up to two mismatches and four mapping sites allowed. Blacklisted genomic regions (https://sites.google.com/site/anshulkundaje/projects/blacklists, (24)) and PCR artifacts were filtered out and peaks were called by using MACS2 (version 2.1.0) with parameter ‐‐*shift* -*100* ‐‐ *ext 200* at significance threshold of FDR 0.01. Data quality was evaluated using standard quality control (QC) metrics: (i) number of uniquely mapped reads, (ii) PCR Bottleneck Coefficient (PBC), (iii) Normalized Strand Cross-correlation coefficient (NSC), and (iv) Relative Strand Cross-correlation coefficient (RSC) (25). QC scores were comparable to those for available ENCODE DNase-seq data (**Table S2**).

Differential DNase-seq peaks were determined using DESeq (26). The top 100k highest confidence DNase-seq peaks from fibroblasts and myoblasts (called by MACS2 as described above) were merged using “bedtools merge” with the default “-d 0” option. Next, DESeq (26) was called on the merged set of DNase I hypersensitive (DHS) regions from the autosomal chromosomes (N=128,080) to identify differentially accessible chromatin sites between the untransfected fibroblasts and the myoblast cells. DNase-seq read counts were computed for each DHS region for each of the three fibroblast and four myoblast replicates available. All DHS regions with at least two-fold differential enrichment and adjusted p-value ≤ 0.01 were selected as differentially accessible sites between fibroblasts and myoblasts. Among these sites, DHS sites with significantly higher signal in fibroblasts were defined as “fibroblast-specific”. Similarly, DHS sites with significantly higher signal in myoblasts were defined as “myoblast-specific”.

### RNA-seq and defining fibroblast- or myoblast-specific genes

RNA was extracted and purified by the methods in RNeasy Mini Handbook from Qiagen. RNA-seq libraries were made by standard TruSeq library preparation with PolyA selection, and processed 50bp paired-end sequencing on Illumina Hi-Seq 2000 platform in the Duke Sequencing and Analysis Core Resource. After trimming bases with low quality score, reads were aligned to the UCSC Genes hg19 reference transcriptome and filtering sequences containing no insertion, using Tophat with options -x 4 and -n 2 (27). Data quality was assessed using RNASeQC (28). All samples showed high mapping rate, low number of alternative aligned reads, and low rate of mismatched bases (**Table S2**). Fragments Per Kilobase of exon per Million fragments mapped (FPKM) was estimated by Cufflinks and differential expression between normalized gene read counts in FPKM with significance threshold of FDR 0.05 by using Cuffdiff (29). To compare the degree of chromatin versus gene expression reprogramming (**Figure S13**), transcript per million (TPM) values were computed for all UCSC known genes in the hg19 assembly.

Given our focus on fibroblast-to-myoblast conversion, we define “fibroblast-specific” genes as genes with significantly higher expression levels in fibroblasts compared to myoblasts, as determined from the RNA-seq data (see above). Similarly, we define “myoblast-specific” genes as genes with significantly higher expression levels in myoblasts compared to fibroblasts (this set includes, but it is not limited to, genes characterized in the literature as being muscle-specific). Using the definitions above, we identified 220 fibroblast-specific genes and 268 myoblast-specific genes.

### ChIP-seq of MyoD-reprogrammed fibroblasts cells

Chromatin immunoprecipitation was processed with two biological replicates of fibroblasts cells following induction of MyoD expression; each replicate contains 20 million cells. One of the biological replicates had two technical replicates. Briefly, cells were fixed with 1% formaldehyde (wt/vol) for 15 min, washed with 1 X PBS and lysed in buffer with 50 mM Tris(pH8), 1% SDS (wt/vol), and 10 mM EDTA. Lysed cells were sonicated with 30 s on/off cycles at high intensity by using a bioruptor (Diagenode). Sonicated supernatants were diluted with buffer containing 16.7 mM Tris-HCl(pH8), 0.01% SDS, 1.1% Triton X-100, 1.2 mM EDTA and 167 mM NaCl. 10 ul of FLAG antibody (Monoclonal anti-FLAG M1 antibody produced in mouse, F3040, Sigma-Aldrich) was added into the diluted supernatants, and 60 ul of Dynabeads protein A beads were added and incubated for 3 hours at 4 °C. ChIP-seq libraries were made by using NEBNext Ultra DNA Library prep kit for Illumina, and 50 bp single-end sequencing was processed on Hi-Seq 2000 platform in the Duke Sequencing and Analysis Core Resource. Sequences were aligned to hg19 reference genome by using bowtie-0.12.9. Uniquely aligned reads were used for peak calling with MACS2 (version 2.1.0) (see below). Data quality was evaluated using standard quality control (QC) metrics: (i) number of uniquely mapped reads, (ii) PCR Bottleneck Coefficient (PBC), (iii) Normalized Strand Cross-correlation coefficient (NSC), and (iv) Relative Strand Cross-correlation coefficient (RSC) (25). QC scores were comparable to those for available ENCODE ChIP-seq data (**Table S2**).

### Calling peaks from MyoD ChIP-seq data

MACS2 v2.10 (30) was used to call peaks for each replicate, and the fragment length input was estimated based on the strand cross-correlation plot generated using SPP (31). For the MyoD ChIP-seq data on fibro-MyoD cells, the reproducible peaks were identified using the ENCODE IDR (Irreproducible Discovery Rate) pipeline (32). Specifically, all MyoD peaks with IDR ≤ 0.01 on at least one pair of replicates were selected for downstream analyses. For the MyoD ChIP-seq data on human myoblast cells, MACS2 was run without a control data set, as no control data was available from the study that reported the myoblast ChIP-seq data (33). Given the lack of a control data set, and the fact that only one replicate was available for the myoblast ChIP-seq data, the final set of MyoD peaks in myoblasts were called at stringent FDR threshold of 10e-10.

### Assessing significance of chromatin reprogramming

Given the large number of myoblast-specific DHS sites with Chromatin Reprogramming Level (CRL) scores around 0 (see Equation 1, Results section), we asked whether MyoD induction is in fact leading to significant chromatin remodeling. For this purpose, we used fibro-control DNase-seq data to compute a null distribution of CRL scores. Briefly, for each differentially accessible DHS site *s*, we computed the score *CRL*_*fibro*-*control*_(*s*) = (*DNase*_*fibro*-*control*_(*s*) - *DNase_fibroblast_*(*s*)) / (*DNase_myoblast_*(*s*) - *DNase_fibroblast_*(*s*)), which reflects the level of chromatin accessibility changing due simply to transduction of the human *MYOD1* gene, prior to its induction. As expected based on the fact fibro-control cells are very similar to fibroblasts, we found that the control distributions of *CRL*_*fibro*-*control*_ values are tightly centered around zero (**Figure S4**). In addition, the distributions of CRL scores are significantly different between fibro-MyoD and fibro-control (Wilcoxon signed-rank test p-values < 10e-293, **Figure S4**), indicating that MyoD overexpression does lead to a highly significant level of genome-wide chromatin reprogramming. Nevertheless, many DHS sites failed to be reprogrammed upon MyoD induction.

### Gene ontology enrichment analyses for genes and DHS sites

For the GO enrichment analyses of the fibroblast- or myoblast-specific reprogrammed or non-reprogrammed genes (**Table S3**), the DAVID tool (34,35) was used with the whole genome as background. For the fibroblast-specific or myoblast-specific DHS sites, the GO enrichment analyses were performed using GREAT (36) (**Table S4**) using the whole genome as background and the default “basal plus extension” association rule.

### Clustering TFs based on their DNA-binding specificities

TF-DNA binding specificity features were derived from protein-binding microarray (PBM) data (37). All PBM data sets for mammalian TFs were downloaded from UniPROBE (38). Additionally, DREAM challenge PBM data sets (39) of good quality (defined as data sets for which the derived DNA motifs have at least three consecutive positions with an Information Content ≥ 0.3) were also included. The PBM data sets were used to generate a Pearson correlation coefficient (PCC) matrix. For each pair of data sets, we computed the PCC using only 8-mes with PBM E-score ≥ 0.35 according to at least one of the data sets (i.e. 8-mers specifically bound by the TFs (37)) (**Figure S7**). We then searched for a threshold value *t* such that 90% of all pairs of replicate TF data sets have PCC > *t*. The dendrogram generated from the PCC matrix was then cut at distance corresponding to the PCC threshold, i.e. at a distance *d*=(1-*t*)/2, where *t*=-0.0259. For every cluster, the TF with the smallest average intra-cluster distance was chosen to represent the cluster. The distance between any two TFs was computed as *d*=(1-PCC)/2, where PCC is the Pearson correlation coefficient computed as discussed above. If there were only two elements in the cluster, one was chosen at random. This resulted in 140 clusters, with an average cluster size of 4. A full list of the 140 TF cluster representatives with their corresponding TF cluster members is available in **Table S5**.

### Classification analyses of reprogrammed versus non-reprogrammed DHS sites

We performed classification analyses for the top 1000 reprogrammed versus non-reprogrammed DHS sites, ignoring outliers with very low or very high CRL scores. Specifically, sites with CRL scores lower than -0.1 or higher than 1.1 were considered outliers. Within the [-0.1,+1.1] interval for myoblast-specific sites, the top 1000 sites (i.e. the sites in the “reprogrammed” set) had CRL scores between +0.55 and +1.10, while the bottom 1000 sites (i.e. the sites in the “non-reprogrammed” set) had CRL scores between -0.10 and +0.01. Within the [-0.1,+1.1] interval for fibroblast-specific sites, the top 1000 sites (i.e. the sites in the “reprogrammed” set) had CRL scores between +0.67 and +1.10, while the bottom 1000 sites (i.e. the sites in the “non-reprogrammed” set) had CRL scores between -0.10 and +0.31. We note that the results of our classification analyses are robust to variations in the precise CRL score intervals used to select reprogrammed and non-reprogrammed sites. Before performing the classification, we observed that the GC content was significantly different between the two sets (**Figure S8**), which could result in biased selection of TFs with GC-rich motifs as highly predictive features. In order to correct for the GC bias, we selected subsets of reprogrammed and non-reprogrammed DHS sites that are matched in their GC content, computed in the +/-150bp regions around DHS centers. For each of the 1000 reprogrammed DHS sites, we search for a ‘matching’ non-reprogrammed site with similar GC content (i.e. GC content different by at most 1%). If a matching site was found, then both the reprogrammed DHS site and its matching non-reprogrammed site were selected. Otherwise, the reprogrammed DHS site was filtered out. Non-reprogrammed sites were selected without replacement. This resulted in a total of 847 reprogrammed and 847 non-reprogrammed myoblast-specific DHS sites. Similarly, a total of 531 reprogrammed and 531 non-reprogrammed fibroblast-specific DHS sites were selected for downstream classification analyses.

The following features were used in classification analyses: (1) 140 TF binding specificity features, defined for each TF cluster representative as the maximum 8-mer PBM enrichment score (E-score) in +/- 150bp genomic regions centered at DHS sites, (2) 11 histone modification ChIP-seq signals from the Normal Human Dermal Fibroblasts (NHDF) cell line, computed as read pileups in +/- 0.7kb region around the DHS centers, (3) maximum CTCF and EZH2 ChIP-seq pileup signals in +/- 0.7kb region around DHS centers, obtained by scanning 200bp windows that overlap by 100bp, and (4) normalized fibroblast DNase-seq reads (as computed by DESeq) in the +/- 150bp region around DHS centers.

R packages “randomForest” and “glmnet” were used to run the Random Forest (RF) and Elastic Net (EN) analyses, respectively. RF and EN were repeated 10 times to assess the stability of the classification accuracy and of the feature importance scores, using 75% of the data (randomly selected) for training and the remaining 25% for testing. For RF analyses, the number of trees (ntrees) and number of predictor variables for splitting a node (mtry) were set to their default values. For EN analyses, the best α parameter was selected from the training data using cross-validation by the “cv.glmnet()” function.

### DHS pattern enrichment analysis

The top 100,000 DNase-seq peaks from each of the three cell lines (fibroblast, fibro-MyoD and myoblast) were merged using the “bedtools merge -d 0” command, resulting in a total of 133,714 autosomal merged DHS sites, which were used in subsequent analyses. For every merged DHS site, a binary tag was assigned to indicate whether the DHS site was present (‘1’) or absent (‘0’) in each of the three cell lines. Hence, every merged DHS site was annotated with one of seven binary triplets (‘001’, ‘010’, ‘011’, ‘100’, ‘101’, ‘110’, ‘111’), where the binary tags are for fibroblast, fibro-MyoD and myoblast, in this order.

Next, we checked whether these patterns are significantly enriched or depleted around the genes corresponding to fibroblast-specific or myoblast-specific reprogrammed or non-reprogrammed gene sets, hereby referred to as ‘treatment gene-sets’ (TS), when compared against a corresponding background set (BG). We analyzed four treatment sets: 1) myoblast-specific reprogrammed genes, 2) myoblast-specific non-reprogrammed genes, 3) fibroblast-specific reprogrammed genes, and 4) fibroblast-specific non-reprogrammed genes. For the myoblast-specific treatment sets, the background set was defined as all genes expressed in myoblasts at FPKM >=5. Similarly, for the fibroblast-specific treatment sets, the background set was defined as all genes expressed in fibroblasts at FPKM >=5.

The GSEA algorithm (40) was implemented to perform enrichment analyses for the DHS patterns. For any DHS pattern *p* and gene set TS, all the genes in the corresponding background set BG were sorted in decreasing order of the number of occurrences of pattern *p* in +/- 100kb of the TSS. Then, a running sum statistic, or enrichment score (ES) was computed at every position in the ranked list of genes, as follows:

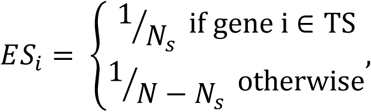

where *N* represents the size of BG and *N_s_* represents the size of TS. The maximum running sum statistic (MES = maximum enrichment score) was recorded as the enrichment for pattern *p* in TS compared to BG. To assess the statistical significance of MES scores, null distributions were generated by randomly permuting the genes in BG 2000 times. Empirical p-values were computed from the null distributions, and corrected for multiple hypothesis testing using the Benjamini-Hochberg procedure (41) as implemented in the “statsmodels.sandbox.stats” python module.

### Computing correlations between chromatin and gene expression reprogramming levels (CRLs vs. GRLs)

For fibroblast- and myoblast-specific genes, we asked whether the extent to which gene expression is reprogrammed correlates with the chromatin reprogramming level (CRL) around the gene. To answer this question, we focused on DHS sites in either 5 kb or 2 kb regions centered at the TSSs. Fibroblast-specific DHS sites were considered in analyses of fibroblast-specific genes, and myoblast-specific DHS sites were considered in analyses of myoblast-specific genes. The CRLs and the gene reprogramming levels (GRLs) were computed in log-scale as follows: CRL(s) or
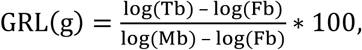
where s and g represent the DHS site and gene under consideration, respectively; Fb, Tb and Mb represent normalized accessibility or transcript per million (TPM) scores for the fibroblasts, transdifferentiated fibroblasts (i.e. fibro-Myod) or myoblasts, respectively. We chose to use log scale in this analysis in order to better observe the entire range of gene expression changes, including small and moderate changes.

## RESULTS

### MyoD induces incomplete genome-wide changes in chromatin accessibility and gene expression

In order to study the extent to which MyoD can transform non-muscle cells into muscle cells, we characterized the genome-wide chromatin accessibility and gene expression profiles of primary human skin fibroblasts, primary human myoblasts, and human fibroblasts that were transdifferentiated by induction of MyoD from a tet-inducible lentiviral vector for 10 days followed by puromycin selection for transduced cells. These cells are henceforth referred to as ‘fibro-MyoD’ (**Figure 1A**). The induction of expression of the *MYOD1* transgene by doxycycline was confirmed by qRT-PCR (**Figure 1B**), western blot (**Figure 1C**), and immunofluorescence using an antibody to MyoD (**Figure S1A**). As expected, no MyoD protein was detected in untransfected fibroblast cells, and minimal MyoD expression was observed in control fibroblast cells transduced with the lentiviral *MYOD1*-containing vector but not induced with doxycycline (cells henceforth referred to as ‘fibro-control’). In fibro-MyoD samples, ~75% of cells showed MyoD-positive nuclei by immunofluorescence staining (**Figure S1A**). In addition, to confirm induction of the *MYOD1* cassette, flow cytometry was used to quantify expression of DsRed, which is co-expressed with MyoD (see Methods) (**Figure S1B**). Transdifferentiation was also confirmed by western blot of myogenin (**Figure 1C**), immunofluorescence staining for skeletal myosin heavy chain (**Figure S1A**), qRT-PCR for additional myogenic markers (**Figure S1C**), and RNA-seq for myogenic markers (**Figure 1D**), similar to previous studies (1,3,13).

**Figure 1.**
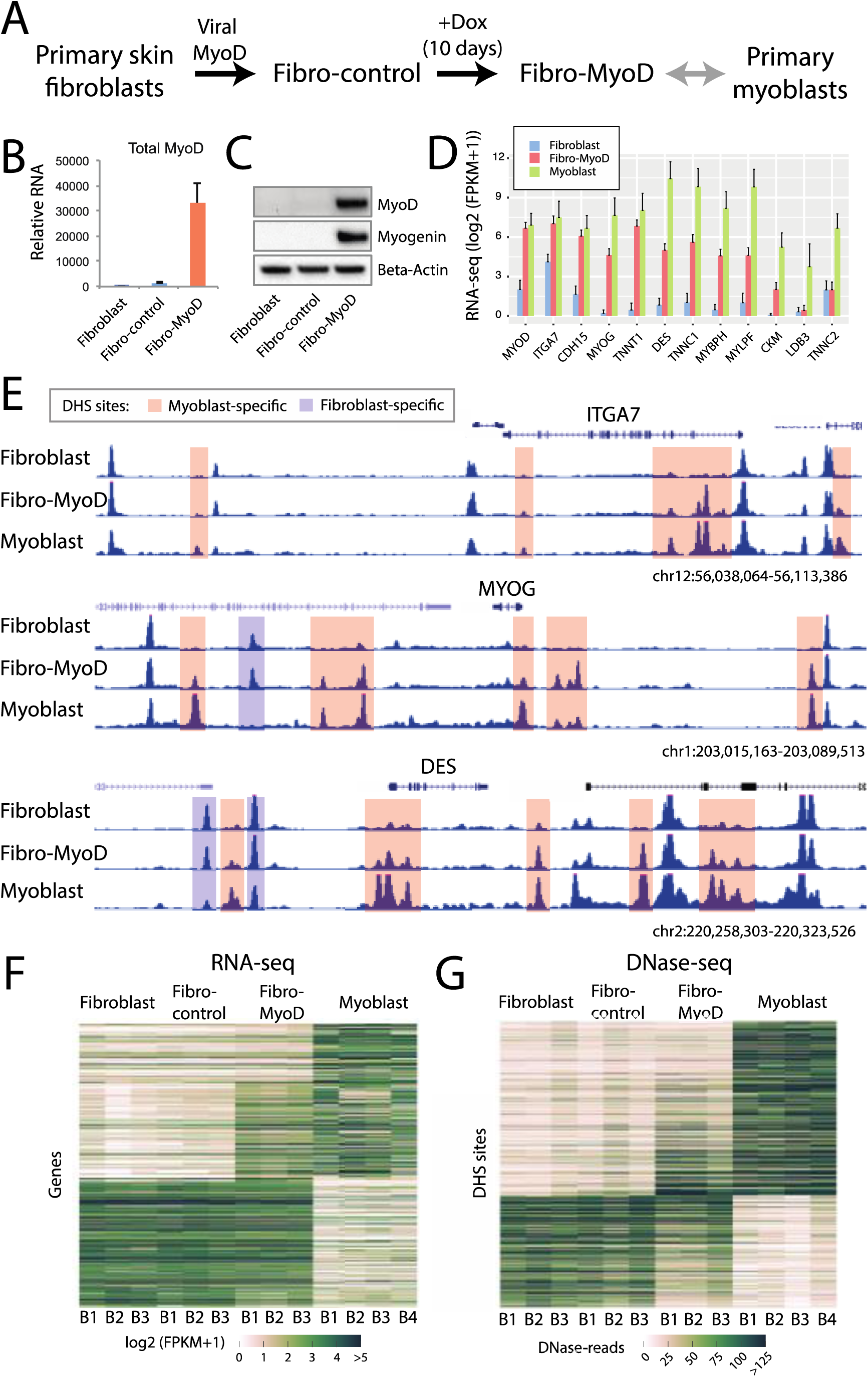
Induced expression of MyoD in fibroblast cells leads to incomplete genome-wide changes in gene expression and chromatin accessibility. **(A)** Schematic representation of the MyoD-induced transdifferentiation experiment performed. **(B)** qRT-PCR and **(C)** western blot showing an increased level of MyoD after induction by addition of doxycycline (dox) for 10 days. Neither MyoD nor another myogenic regulatory factor, Myogenin, is detected in either untransfected fibroblasts or in fibroblasts that contain the MyoD vector but are not treated with doxycycline. **(D)** RNA-seq data (logarithm of fragments per kilobase per million base shifted by 1, or log(FPKM+1)) showing that myogenic genes are significantly increased in fibro-MyoD cells (^**^ = Welch’s t-test p-value < 0.01, ^*^ = Welch’s t-test p-value < 0.05). Error bars show the upper bound of the 95% confidence interval, as computed by Cufflinks. Note that early myogenic genes (ITGA7, CDH15) are reprogrammed, while middle (MYOG) and late (TNNT1, DES, TNNC1, MYBPH, MYLPF, CKM, LDB3, TNNC2) muscle markers are generally partially reprogrammed. **(E)** Representative DNase-seq signal around myogenic genes ITGA7 (early myogenic gene), MYOG (intermediate myogenic gene), and DES (late myogenic gene). Red boxes represent myoblast-specific DHS sites that are partially, completely or not reprogrammed in fibro-MyoD cells. Blue boxes represent fibroblast-specific DHS sites that are maintained in fibro-MyoD cells. Heatmaps of **(F)** RNA-seq and **(G)** DNase-seq for each replicate (B1-B4) of fibroblast, fibro-control, fibro-MyoD, and primary myoblasts are shown for differentially expressed genes and differentially accessible DHS sites, respectively. The data indicate that fibro-Myod cells display partial characteristics of both fibroblasts and myoblasts.

To characterize the genome-wide chromatin changes induced by MyoD, we used DNase-seq to measure chromatin accessibility in fibroblast, fibro-control, fibro-MyoD, and primary myoblast cells (see Methods). We found that for many muscle-specific genes, such as *ITGA7* (integrin, alpha 7), *MYOG* (myogenin) and *DES* (desmin), chromatin accessibility changes in and around these genes are readily detected as a result of MyoD induction, and these changes largely mirror the chromatin accessibility profiles for primary myoblasts (**Figure 1E**). This indicates that the overexpression of MyoD induces chromatin structure changes in a non-random manner that is highly similar to primary muscle cells. We also found that not all myoblast-specific DHS sites (defined here as DHS sites with significantly higher DNase-seq signal in myoblasts compared with fibroblasts; see Methods) open up in fibro-MyoD, with some opening up only partially (**Figure 1E, Figure S2**). This suggests that induction of MyoD alone cannot reorganize chromatin at all potentially myogenic DHS sites in transduced primary human fibroblasts. Furthermore, we also detected fibroblast-specific DHS sites that are not lost after induction of MyoD, indicating that some fibroblast-related regulatory element memory is maintained during the transdifferentiation process (**Figure 1E, Figure S2)**.

This incomplete reprogramming at the chromatin level is reflected in our RNA-seq data, which shows that although the levels of key myogenic marker genes are significantly higher in fibro-MyoD compared to fibroblasts (consistent with our qRT-PCR results), several of these genes are not expressed at the same levels as observed in primary myoblasts (**Figure 1D**). Some of the common myogenic marker genes have been previously designated as important in early (*ITGA7*, *CDH15*), intermediate (*MYOG*), or late (*TNNT1*, *DES*, *TNNC1*, *MYBPH*, *MYLPF*, *CKM*, *LDB3*, *TNNC2*) myogenic differentiation (6,7,11). Our RNA-seq data indicate that ‘early’ genes display a large degree of reprogramming (**Figure 1D**). This result, combined with the majority of transdifferentiated cell nuclei being MyoD-positive (**Figure S1A**), indicates that MyoD induction is indeed differentiating cells toward a myogenic fate. In contrast to ‘early’ genes, ‘late’ marker genes show a wide range of reprogramming levels, with a majority being moderately upregulated and a few showing significant (*TNNC1*) or no (*LBD3* and *TNNC2*) change in expression (**Figure 1D**). It is possible that ‘late’ myogenic genes were not expressed at very high levels in fibro-MyoD cells because transdifferentiation was performed in growth media, without inducing terminal differentiation (see Methods). Nevertheless, according to our direct comparison to myoblasts (which are also not terminally differentiated), some ‘late’ marker genes are, at best, only partially reprogrammed. Importantly, muscle marker genes previously reported to be upregulated during induction of MyoD expression in transfected mouse embryonic fibroblasts incubated in growth medium are also upregulated in our study (6).

In addition to incomplete reprogramming at individually assayed myogenic genes, we also observed incomplete reprogramming at the genome-wide level for both gene expression (**Figure 1F**) and chromatin accessibility (**Figure 1G**). This indicates that fibro-MyoD cells exhibit some of the characteristics of both fibroblasts and myoblasts simultaneously. To verify that the fibro-MyoD cell population is not simply a mixture of undifferentiated (fibroblast) and transdifferentiated (myoblast) cells, we performed *in silico* mixing experiments at different ratios of fibroblast to myoblast cells. As expected, we found that the gene expression and chromatin accessibility profiles of mixed populations (**Figure S3**) are substantially different from the profiles we observed for fibro-MyoD cells (**Figure 1F,G**).

**Figure 1F** shows the expression levels of genes significantly differentially expressed between the starting (fibroblast) and target (myoblast) cell types. We refer to these genes as fibroblast-specific or myoblast-specific, depending on whether the genes are significantly more expressed in fibroblasts or myoblasts, respectively (see Methods). As expected, all genes have similar expression levels between fibroblasts and fibro-control cells. After MyoD induction, some genes are up- or down-regulated and, as a result, their expression levels become similar to those in myoblasts – i.e., the genes have their expression reprogrammed. However, a large fraction of fibroblast-specific or myoblast-specific genes also remain non-reprogrammed. We identified these reprogrammed and non-reprogrammed genes through pairwise comparisons of the expression levels in fibroblasts, fibro-MyoD and myoblasts, as computed by Cuffdiff (see Methods). In order to assess the functional significance of these reprogrammed or non-reprogrammed genes, we performed gene ontology (GO) analysis using DAVID (34,35). The myoblast-specific non-reprogrammed genes (N=100), as well as the reprogrammed genes (N=168), are highly enriched for muscle-specific GO categories such as “sarcomere” and “striated muscle contraction” (**Table S3**). This indicates that non-reprogrammed genes are also relevant to muscle biology. Similarly, both reprogrammed (N=34) and non-reprogrammed (N=186) fibroblast-specific genes are enriched for fibroblast-associated terms such as “extracellular matrix structural constituent” (**Table S3**). These observations suggest that, based on its gene expression profile, the fibro-MyoD cell line represents an intermediate between fibroblast and myogenic cells.

Similar to the gene expression analysis, global chromatin accessibility (DNase-seq) data revealed the fibro-MyoD cell line represents an intermediate between fibroblast and myogenic cells (**Figure 1G**). While many DHS sites are well reprogrammed (i.e. their accessibility level is similar to that in myoblasts, see Methods), a large number of sites that are different between fibroblasts and myoblasts do not change their accessibility profile after MyoD induction. To assess the potential regulatory activities of these sites, we performed GO enrichment analyses using GREAT (36) (**Table S4;** see Methods). As expected, myoblast-specific DHS sites that open up in fibro-MyoD (i.e. are reprogrammed) are highly enriched for GO annotations pertaining to muscle and satellite cell differentiation. In addition, myoblast-specific DHS sites that remain closed in fibro-MyoD (i.e. are non-reprogrammed) also show enrichment for muscle and regeneration ontologies, albeit to a lower extent than for the reprogrammed DHS sites. Similarly, fibroblast-specific DHS sites that lose accessibility (i.e. are reprogrammed) are associated with fibroblast-specific genes that have been previously reported to be down-regulated in response to induction of MyoD expression in transduced 3T3 mouse fibroblasts (42) (**Table S4**). In addition, fibroblast-specific DHS sites that remain open during transdifferentiation (i.e. are not reprogrammed) are involved in epithelial-mesenchymal transition, cell motility and interstitial matrix, which are fibroblast-associated annotations. The fact that we observe such non-reprogrammed fibroblast-specific DHS sites is consistent with previous reprogramming studies that also found evidence of an “epigenetic memory” from the cells of origin (43-45).

### MyoD induction results in a continuum of reprogrammed DHS sites

In order to quantify reprogramming efficiency at DHS sites differentially accessible between fibroblasts and myoblasts, we introduce the ‘chromatin reprogramming level’ (CRL) score. For each differential DHS site *s*, the CRL score is defined as the ratio of the change in chromatin accessibility due to MyoD induction in fibroblasts relative to the difference in accessibility between fibroblasts and myoblasts:

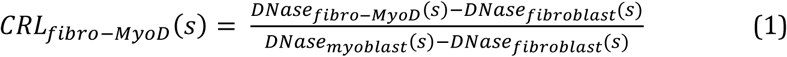

where *DNase_c_*(*s*) represents the normalized DNase-seq read signal at DHS site *s* in cell type *c* (see Methods). For both fibroblast- and myoblast-specific DHS sites, a CRL score close to 0 indicates that the site *s* is not reprogrammed, i.e. its chromatin accessibility in fibro-MyoD cells is similar to that in fibroblasts, while a CRL score close to 1 indicates that the site *s* is reprogrammed, i.e. its chromatin accessibility in fibro-MyoD cells changed significantly and is similar to the level of chromatin accessibility observed in myoblasts (**Figure 2A,B**).

**Figure 2:**
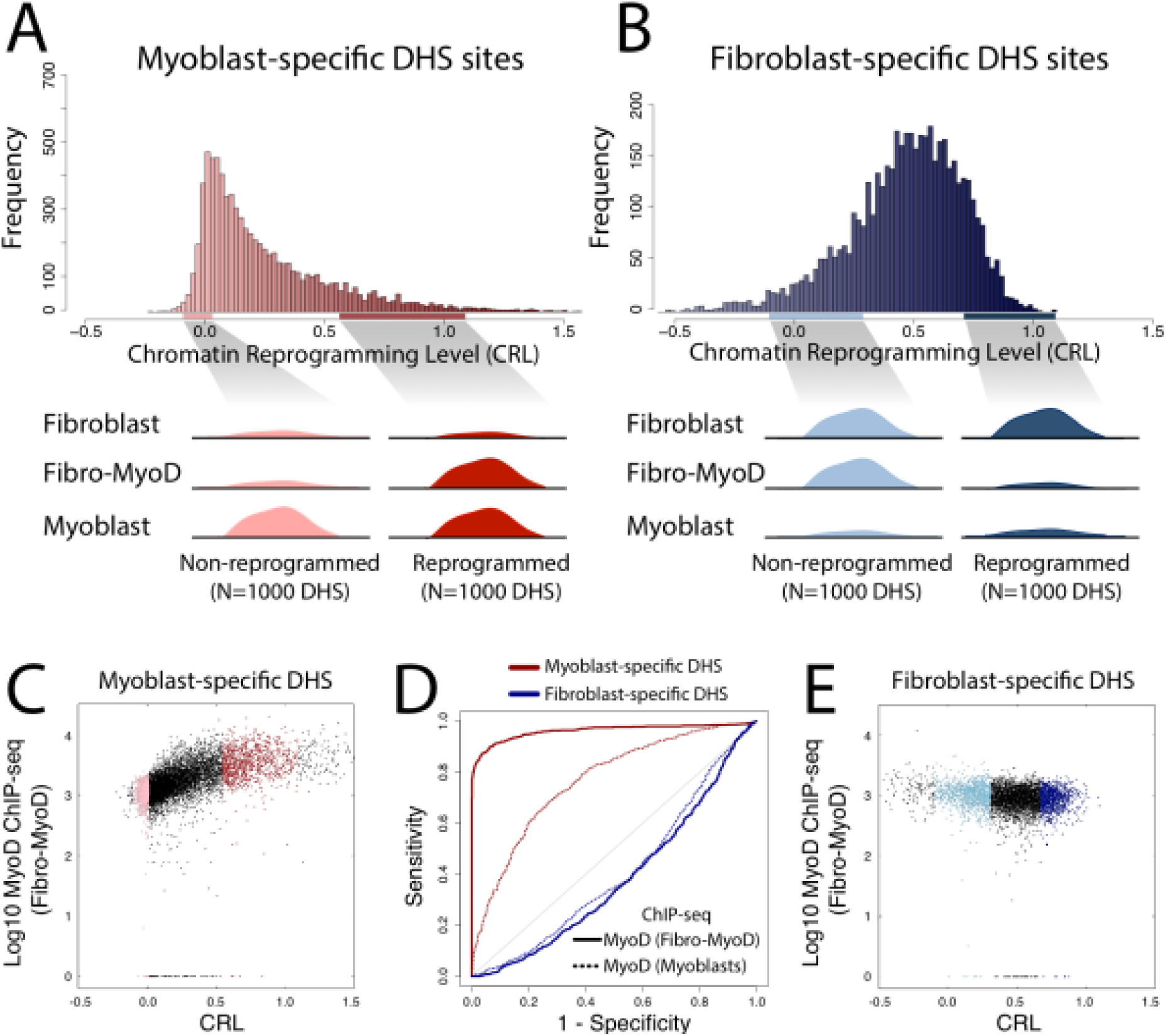
MyoD induction results in continuum of chromatin reprogramming genome-wide. **(A)** Distribution of chromatin reprogramming level (CRL) scores for myoblast-specific DHS sites. For classification analyses we selected 1000 non-reprogrammed sites and 1000 reprogrammed sites from the ends of the distribution, ignoring potential outliers with extreme CRL scores. Schematics of sites that are not reprogrammed (pink) or reprogrammed (dark red) are shown. **(B)** Analogous to panel A but for fibroblast-specific DHS sites. **(C)** Scatterplot showing positive correlation between CRL scores and MyoD ChIP-seq signal for myoblast-specific DHS in fibro-MyoD cells (Spearman correlation coefficient: 0.66). The colors for the top 1000 reprogrammed or non-reprogrammed sites (dark red and pink, respectively) correspond to panel A. **(D)** ROC curves for classification of reprogrammed versus non-reprogrammed DHS sites that are myoblast-specific (red) or fibroblast-specific (blue), obtained using MyoD ChIP-seq signal from either fibro-MyoD cells (solid lines) or from myoblasts (dotted lines). **(E)** Analogous to panel C, but for fibroblast-specific DHS sites (Spearman correlation coefficient: -0.17).

We detect a continuum of CRL scores for myoblast-specific (**Figure 2A**) and fibroblast-specific (**Figure 2B**) DHS sites that span mostly between 0 (non-reprogrammed sites) and 1 (reprogrammed sites). Interestingly, while CRL distribution for myoblast-specific DHS sites is positively skewed (median of only 0.16), the median CRL for fibroblast-specific DHS sites is 0.49. This suggests that reprogramming efficiency is generally higher for fibroblast-specific compared to myoblast-specific sites. We confirmed that these positively skewed CRL scores are significant by comparing them to CRL scores computed using fibro-control DNase-seq data (**Figure S4** and Methods).

In order to understand why many DHS sites fail to open up during transdifferentiation, we first compared two sets of myoblast-specific sites: a set containing 1000 non-reprogrammed sites with the smallest CRL scores, and another set containing 1000 reprogrammed sites with the largest CRL scores, ignoring potential outliers (**Figure 2A**, see Methods). Similarly, in order to understand why some DHS sites fail to close down during transdifferentiation, we compared two sets of 1000 fibroblast-specific DHS sites with CRL values indicative of reprogrammed versus non-reprogrammed status (**Figure 2B**, see Methods). Using the control distribution of CRL scores derived from fibro-control DNase-seq data (**Figure S4**), we estimate the false discovery rate (FDR) for the 1000 reprogrammed DHS sites to be 0.01 and 0.005 for myoblast-specific and fibroblast-specific sites, respectively (see Methods). Due to a low level of leaky expression of MyoD in fibro-control cells (**Figure 1B**), these FDR values are likely conservative. Importantly, the results presented below are robust to variations in the precise ranges of CRL scores used to call the 1000 reprogrammed and 1000 non-reprogrammed sites.

Previous small-scale studies reported that MyoD can open chromatin and attract chromatin-remodeling enzymes to myogenic enhancers (18,42,46). Our ChIP-seq data for MyoD in fibro-MyoD cells shows that about half of the binding sites occurred in previously open chromatin regions, while the other half occur at sites that were closed in fibroblast cells but opened up in response to MyoD overexpression. We also found that the majority of these MyoD binding sites in fibro-MyoD are also accessible in myoblasts (**Figure S5**). Therefore, we asked whether the lack of chromatin reprogramming at myoblast-specific DHS sites might be correlated with lack of MyoD binding. Indeed, we observed a positive correlation between the MyoD ChIP-seq signal in fibro-myoD cells (quantified at +/- 150bp of DHS center) and the corresponding CRL values (Spearman correlation coefficient: 0.66, **Figure 2C**). Moreover, we also found that the MyoD ChIP-seq signal from fibro-MyoD cells can almost perfectly separate reprogrammed from non-reprogrammed myoblast-specific DHS sites (area under the receiver-operating characteristic curve (AUC): 0.96, **Figure 2D**, solid red curve). Broadly, these results indicate that MyoD binding in transdifferentiated cells is strongly associated with chromatin opening, while non-reprogrammed myoblast-specific sites lack MyoD binding. We performed a similar classification analysis using the MyoD ChIP-seq signal from primary myoblasts (33), and found that binding of MyoD in myoblasts is less accurate in distinguishing reprogrammed from non-reprogrammed DHS sites (AUC: 0.76, **Figure 2D**, dotted red curve). This drop in AUC is likely due to the fact that ~33% of myoblast-specific DHS sites that are not reprogrammed and not bound by MyoD in transdifferentiated cells show MyoD ChIP-seq peaks in myoblasts (at MACS2 FDR cutoff 1e-10, see Methods). Overall, these results show a large but not complete overlap between MyoD-bound sites in fibro-MyoD and myoblasts (**Figure S6**).

In contrast to MyoD being closely associated with the opening of chromatin at myoblast-specific DHS sites, MyoD binding in fibro-MyoD cells is slightly inversely correlated with chromatin closing at fibroblast-specific sites (Spearman correlation coefficient: -0.17, **Figure 2E**). Hence, MyoD binding is a weak predictor of reprogramming status of fibroblast-specific sites (AUC ~0.38, **Figure 2D**). This suggests that binding of MyoD at some already open chromatin sites could potentially prevent closing down of these sites, but this would only explain a small fraction of non-reprogrammed sites. Indeed, only 8 of the 1000 non-reprogrammed fibroblast-specific sites show MyoD binding signal in fibro-MyoD (ChIP-seq peaks called by MACS2 and filtered at IDR 0.01, see Methods).

### TF motifs distinguish reprogrammed from non-reprogrammed myoblast-specific DHS sites

To understand why the non-reprogrammed DHS sites are not bound by MyoD in fibro-MyoD cells, and what other factors might contribute to their lack of MyoD binding and/or their lack of reprogramming, we used a classification approach to distinguish the top 1000 reprogrammed sites from the top 1000 non-reprogrammed sites based on genomic and epigenomic features. The features used for this analysis were: 1) how open the site was in the starting cell type (i.e. its DNase-seq signal in fibroblast cells); 2) the pre-existing chromatin environment at the DHS site, as reflected by ChIP-seq signal for 11 histone marks, CTCF, and EZH2 proteins in normal human dermal fibroblast (NHDF) cells (24); and 3) the DNA binding specificities of various TFs for the DHS site, as determined using protein-binding microarray (PBM) data (see Methods) (37). To derive TF-DNA binding specificity features, we used PBM data for human and mouse TFs (38,39). In addition, since PBM data are not available for human MyoD, we used data for the MyoD ortholog in *C. elegans*, HLH-1, henceforth referred to as cMyoD. Since many TFs in our data set have similar DNA binding specificities, which could lead to high collinearity among features used in our classifier, we clustered the PBM data into 140 clusters (Methods). For each cluster, we selected a representative TF that minimizes the intra-cluster distance (**Table S5**; Methods), and we used as a feature the maximum DNA binding specificity score (i.e. the maximum 8-mer PBM E-score) in the +/- 150bp genomic region around the center of the DHS site (see Methods). The PBM E-score is a modified form of the Wilcoxon-Mann Whitney statistic and ranges from -0.5 (least favored sequence) to +0.5 (most favored sequence), with values above 0.35 corresponding, in general, to sequence-specific TF-DNA binding (37) (**Figure S7**).

To ensure that the two sets of sequences used in our classification have similar GC content, we randomly sampled reprogrammed and non-reprogrammed sites so that the GC-content distributions for the two sets of sites were matched (**Figure S8**). This reduced the number of sites in each class from 1000 to 847. This step was necessary to avoid the potential bias of selecting certain TFs as predictive features simply because of the GC-content of their motifs. Next, to perform the classification, we randomly selected 75% of the input sequences to train a Random Forest or an Elastic Net model, and used the remaining 25% for testing. We repeated this training-testing step 10 times in order to get an assessment of the stability of the classification accuracy, as well as the stability of the feature ranking. The results of our Random Forest classification analysis (**Figure 3**) are similar to the results obtained using Elastic Net (**Table S6**) and also similar to the results obtained for sites not matched by GC content (**Figure S9**).

**Figure 3:**
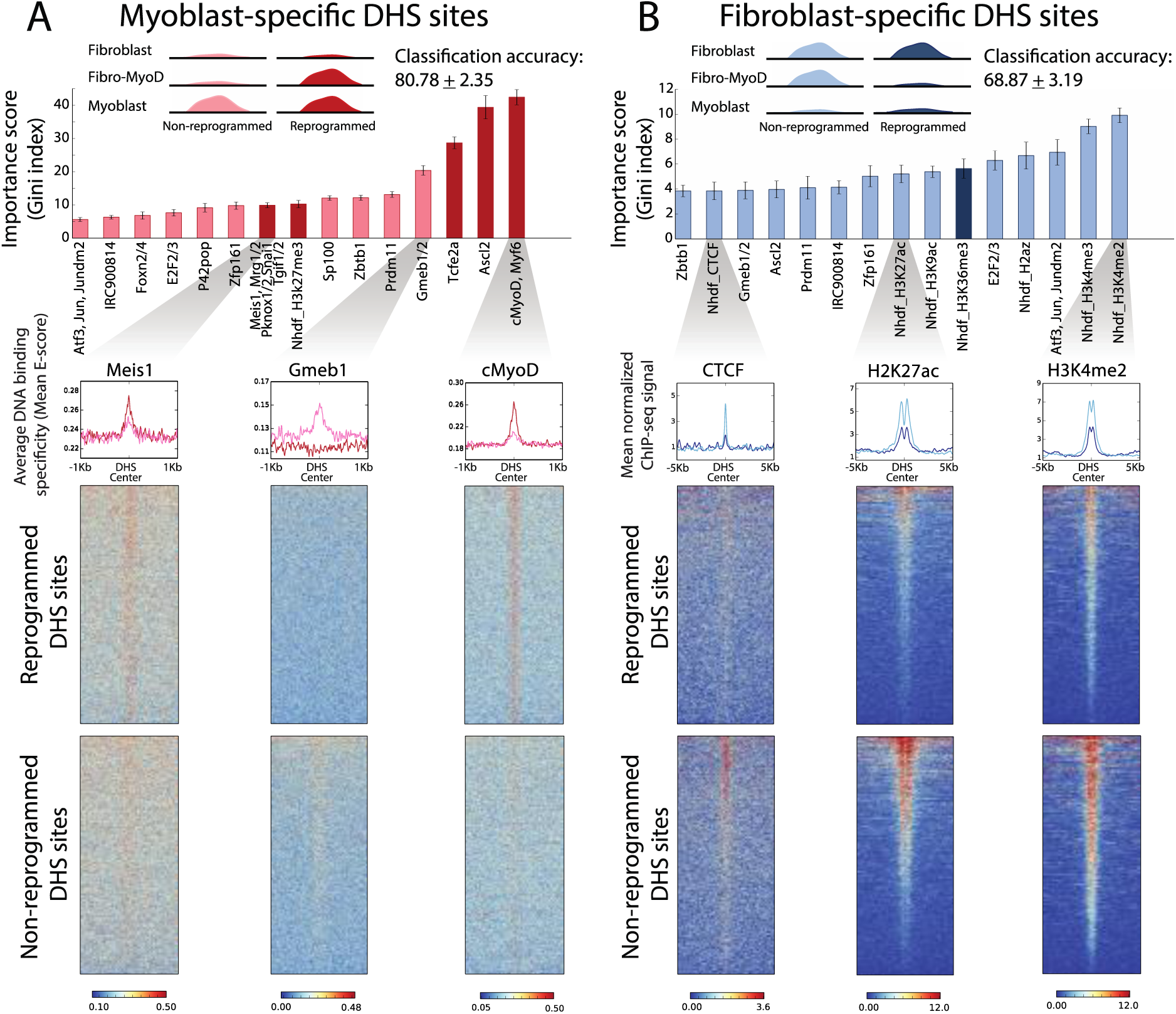
Genetic and epigenetic features that distinguish reprogrammed from non-reprogrammed DHS sites. **(A)** For myoblast-specific sites, the reprogramming status is largely predicted by DNA binding specificities of MyoD, MyoD cofactors (Tcfe2a, Meis1/Pknox1), and SAND-domain factors (Gmeb1/2, Sp100). **(B)** For fibroblast-specific sites, the most predictive features include several active epigenetic marks (H3K4me2, H3K4me3, etc.) that were present before MyoD induction. Top panels show the classification accuracy and the top 15 most predictive features obtained from Random Forest classifiers. For each feature, the color of the bar corresponds to the class most correlated with the feature (reprogrammed = dark color; non-reprogrammed = light color). Bottom panels show, for selected features, the heatmaps corresponding to either the DNA binding specificities of the TF cluster representatives (in panel A), or the ChIP-seq signal for histone marks and CTCF in a dermal fibroblast cell line, NHDF (in panel B). The summary plots above each heatmap show the average signal for either of the two classes. The heatmaps and summary plots were generated using deepTools (72).

Using a Random Forest classifier on reprogrammed versus non-reprogrammed sites, we obtained a high classification accuracy of 80.78 +/- 2.35% (**Figure 3A**). The most predictive feature for distinguishing reprogrammed from non-reprogrammed sites is the DNA-binding specificity of cMyoD and Myf6, which are myogenic regulatory factors (MRFs) with similar DNA binding specificity (47). This is consistent with our results in **Figure 2D**. The greater binding specificity of cMyoD/Myf6 for DNA sequences present in reprogrammed vs. non-reprogrammed DHS sites indicates that the lack of MyoD binding at non-reprogrammed DHS sites can be largely attributed to low MyoD-DNA binding affinity. The second most predictive feature is the DNA binding specificity of ASCL2, which is a basic helix-loop-helix factor involved in neuronal differentiation that has a DNA binding specificity similar to MyoD (48-50). The third most predictive feature is the DNA-binding specificity of TCFE2A (or E2A), a mouse E-protein that is homologous to human E12/E47, known heterodimer partners of MyoD (51). Our classification models also selected other known cofactors of MyoD, namely PKNOX1/MEIS1, as features predictive of chromatin opening (52-55). Thus, reprogrammed myoblast-specific DHS sites are largely characterized by a high affinity for MyoD and its co-factors. In contrast, the non-reprogrammed myoblast-specific DHS sites have lower affinity for MyoD and its cofactors, but higher affinity for SAND-domain factors such as GMEB1, GMEB2, and SP100. SAND-domain proteins, which are a class of DNA-binding proteins involved in chromatin-associated transcriptional regulation (56), were only minimally expressed in fibro-MyoD (**Table S7**).

Compared to the TF features, epigenetic features based on fibroblast histone modification data played only a minor role in the classification of myoblast-specific sites. The most predictive epigenetic mark, H3K27me3, was moderately enriched at reprogrammed sites compared to non-reprogrammed sites (**Figure S10**). This pre-existing relative abundance of H3K27me3 at MyoD-induced reprogrammed sites is in agreement with similar observations at genomic sites targeted by other pioneering factors, such as NEUROD1 during neuronal differentiation (57) and PU.1 during remodeling induced by histone deacetylase inhibitors (HDACi) (58). It is known that H3K27me3, when concomitantly observed with H3K4me1 or H3K4me2, marks poised enhancers (59). Because we also see pre-existing H3K4me1 and H3K4me2 enrichment at myoblast-specific reprogrammed chromatin sites, albeit non-differential from non-reprogrammed sites (**Figure S10**), this suggests that MyoD is potentially targeting poised enhancers during reprogramming.

### Histone modification marks distinguish reprogrammed from non-reprogrammed fibroblast-specific DHS sites

We performed similar classification analyses for reprogrammed versus non-reprogrammed fibroblast-specific DHS sites (**Figure 3B**), using the same set of features described above. As for myoblast-specific sites, the GC-content was matched between reprogrammed and non-reprogrammed fibroblast-specific DHS sites (**Figure S8**), resulting in 531 sequences in each class. The classification accuracy achieved by the Random Forest classifier on fibroblast-specific sites was 68.87 +/- 3.19% (**Figure 3B**). Similar to myoblast-specific sites, the classification accuracies and predictive features selected by Elastic Net models largely match those of Random Forest models (**Table S6**).

Unlike myoblast-specific sites, the majority of the top predictors for fibroblast-specific site classification are histone marks. H3K4me2, H3K4me3, H3K27ac, H2az, and H3K9ac are enriched at non-reprogrammed vs. reprogrammed fibroblast-specific DHS sites in fibroblasts, suggesting that sites with these active marks are refractory to direct or indirect heterochromatization by MyoD. Importantly, this is true despite the fact that the pre-existing chromatin accessibility is not discriminatory between the sets of sites (**Table S6, Figure S11**). In addition, we found that high-affinity sites for the activator proteins E2F2/E2F3 and *in vivo* binding of CTCF in fibroblast cells, were enriched in the non-reprogrammed compared to the reprogrammed sites, suggesting that binding of these factors could be involved in maintaining chromatin accessibility at some sites.

### Chromatin remodeling deficiencies are associated with nearby non-reprogrammed genes

To determine whether chromatin-remodeling deficiencies can explain the incomplete reprogramming observed at the gene level, we tested whether different types of chromatin remodeling events are enriched around reprogrammed and non-reprogrammed genes (**Figure 4**, Methods). DHS sites from fibroblasts, fibro-MyoD and myoblasts were binarized with respect to state of chromatin accessibility (1 = open, 0 = closed). For example, the ‘011’ DHS pattern refers to sites that are closed in fibroblasts (0), open in fibro-MyoD (1), and open in myoblasts (1). Next, for each possible pattern we asked whether it is significantly enriched in the neighbourhood of genes in test sets compared to background sets. We analyzed four test sets: 1) myoblast-specific reprogrammed genes (**Figure 4A**), 2) myoblast-specific non-reprogrammed genes (**Figure 4D**), 3) fibroblast-specific reprogrammed genes (**Figure S12A**), and 4) fibroblast-specific non-reprogrammed genes (**Figure S12C**). For the myoblast-specific treatment sets, the background set was defined as all genes expressed in myoblasts at FPKM ≥ 5. Similarly, for the fibroblast-specific treatment sets, the background set was defined as all genes expressed in fibroblasts at FPKM ≥ 5. To perform enrichment analyses for DHS patterns we implemented the GSEA algorithm (40), sorting genes by the number of occurrences of a pattern in the genomic regions +/- 100kb of gene TSSs (see Methods). We used the Benjamini-Hochberg procedure to correct for multiple hypothesis testing, and characterized DHS patterns enriched at a false discovery rate (q-value) < 0.05.

**Figure 4:**
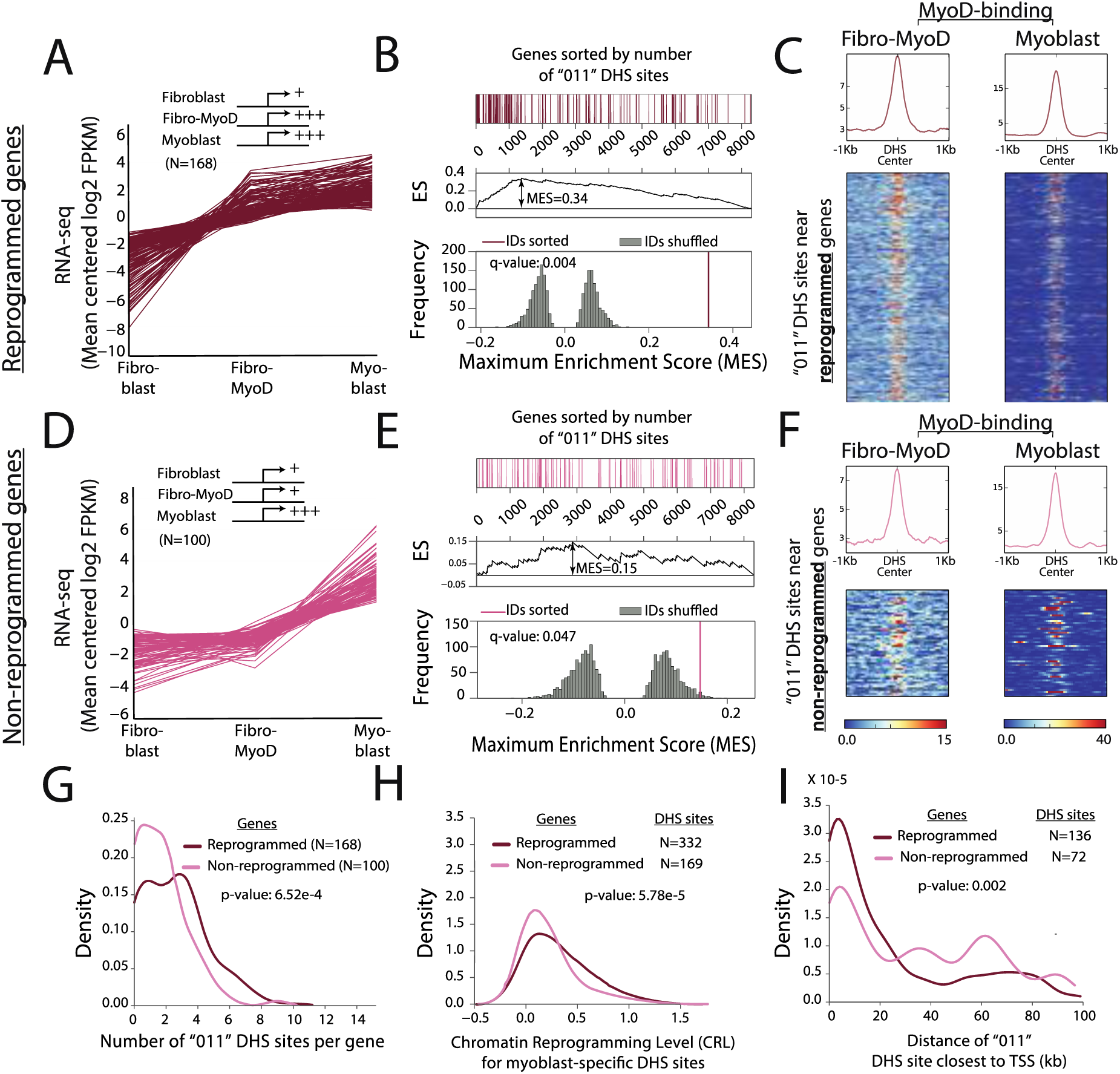
Reprogrammed genes are associated with reprogrammed chromatin profiles. **(A)** Normalized gene expression levels for reprogrammed myoblast-specific genes. Inset shows a schematic representation of the gene expression level in the three cell lines. **(B)** GSEA analysis (40) showing reprogrammed ‘011’ DHS sites are significantly enriched in the +/- 100kb regions around TSSs of reprogrammed genes. Heatmap shows all genes expressed in myoblasts (FPKM >=5) sorted by the number of ‘011’ DHS sites, with reprogrammed genes shown in red and all other genes shown in white. Based on the ordered list of genes, we computed an enrichment score (ES) at every position in the list, and used the maximum enrichment score (MES) as our test statistic. We assessed the significance of the test statistic by comparing it to a null distribution (grey histogram) computed from 1,000 random gene sets (see Methods). **(C)** MyoD binding is observed at reprogrammed ‘011’ DHS sites in both fibro-MyoD and myoblast cells. **(D-F)** Similar to A-C, but for non-programmed myoblast-specific genes. **(G)** Reprogrammed genes have significantly more ‘011’ DHS sites compared to non-reprogrammed genes (one-sided Mann-Whitney U test p-value: 6.52e-4). **(H)** Chromatin reprogramming level (CRL) is higher for DHS sites around reprogrammed compared to non-reprogrammed genes (one-sided Mann-Whitney U test p-value: 5.78e-5). **(I)** Reprogrammed genes have ‘011’ DHS sites closer to the TSS compared to non-reprogrammed genes (one-sided Mann Whitney U test p-value: 2e-3). For each gene, we considered the ‘011’ DHS site closest to its TSS, if present within 100kb of the TSS.

Reprogrammed (‘011’) DHS sites are enriched around reprogrammed genes (**Figure 4B**, q-value: 0.004), which indicates that these sites may be bound by regulatory factors including MyoD (**Figure 4C**) required for turning on myoblast-specific genes. We also found that ‘011’ DHS sites are modestly enriched around non-reprogrammed genes (q-value: 0.047, **Figure 4E**), suggesting that some myogenic regulatory regions that open up in response to MyoD induction could still be missing critical trans factors necessary to activate gene expression. Classification of ‘011’ DHS sites nearby reprogrammed versus non-reprogrammed genes, using the same genetic and epigenetic features mentioned above, was not better than random, possibly also indicating missing non-DNA binding co-factors which our classification cannot identify. In addition, we found that ‘011’ sites around reprogrammed versus non-reprogrammed genes have similar levels of MyoD binding in fibro-MyoD (Figures 4C and 4F, left panels) and myoblast cells (**Figures 4C** and **4F**, right panels).

Although both reprogrammed and non-reprogrammed gene sets show over-representation of ‘011’ DHS sites, there exist significant differences between the two sets themselves. First, reprogrammed vs. non-reprogrammed genes have significantly more ‘011’ DHS sites in their neighborhood (within 100kb; p-value: 6.52e-4, **Figure 4G**), which suggests that although non-reprogrammed genes have chromatin opening more than one would expect by chance (i.e. more than control gene sets), they may still lack sufficient chromatin reprogramming. Second, compared to non-reprogrammed genes, the reprogrammed genes displayed a significantly higher degree of chromatin opening (p-value: 5.78e-5, **Figure 4H**). Third, chromatin reprogramming occurred closer to the TSS for reprogrammed myoblast-specific genes (p-value: 2e-3, **Figure 4I**).

We observed similar results for fibroblast-specific genes (**Figure S12**). Both the reprogrammed and non-reprogrammed gene sets are enriched for reprogrammed (‘100’) and non-reprogrammed (‘110’) DHS sites (q-values < 0.006; Methods). The abundance of reprogrammed ‘100’ sites is not different in the two sets of genes (**Figure S12E**), but the CRL scores (i.e. levels of chromatin reprogramming) are significantly higher around reprogrammed genes (one-sided Mann-Whitney U test p-value: 0.005, **Figure S12F**). Interestingly, non-reprogrammed (‘110’) chromatin sites are significantly closer to the TSSs of non-reprogrammed versus reprogrammed fibroblast-specific genes (one-sided Mann-Whitney U test p-value: 0.006, **Figure S12G**), indicative of an association between gene expression and promoter-region accessibility.

We also assessed whether promoter-proximal chromatin remodeling, or lack thereof, explains the variation in gene expression reprogramming (**Figure S13**). For this analysis, we focused on promoter-proximal (TSS ± 5kb or TSS ± 2kb) myoblast-specific and fibroblast-specific DHS sites (Methods). The extent of gene expression reprogramming level (GRL) was computed similarly to CRL (Methods). Log-transformed values were used to compute both CRLs and GRLs, in order to capture even small and moderate changes in expression level and accessibility. We observed strong correlation between the degree of chromatin reprogramming and the degree to which gene-expression is reprogrammed (PCC: 0.48, p=2.7e-16 for myoblast-specific genes; PCC: 0.41, p=0.0044 for fibroblast-specific genes at TSS ± 5kb, **Figure S13A**). The observed correlation was stronger for TSS ± 2kb regions (PCC: 0.6, p=6.4e-18 for myoblast-specific genes; PCC:0.48, p=0.0079 for fibroblast-specific genes, **Figure S13B**), suggesting that chromatin accessibility remodeling in promoter regions partially explains the variation in reprogramming of gene expression during MyoD-induced transdifferentiation. To our knowledge, this is the first genome-wide study comparing gene expression and chromatin accessibility profiles of myoblasts versus MyoD-transdifferentiated cells.

## DISCUSSION

The MyoD-induced transdifferentiation model system can be used to understand how master regulatory TFs can transform cell fates by inducing global changes in chromatin and gene expression profiles. However, it is unknown at a genome-wide scale how much transdifferentiated cells quantitatively differ from both the starting cells and the target cells. Here, we use MyoD-induced transdifferentiation of primary human skin fibroblasts to the myogenic cell fate as a model system to develop a general approach for investigating the extent of chromatin- and gene-level reprogramming induced by forced overexpression of TF master regulators. In our system, we find that while many of the early muscle marker genes are reprogrammed, global gene expression and accessibility changes are still incomplete when compared to myoblasts, the early myogenic determination stage. Our findings suggest how incomplete transdifferentiation can be quantified, characterized, and potentially improved.

This study is the first to quantify the effects of MyoD on chromatin accessibility in a transdifferentiation system. We are using primary human cells, which are known be more challenging to reprogram with MyoD compared to murine and/or immortalized cell lines (1,4,5,7,12,14,19,60,61). In this system, we found that MyoD targets both closed and accessible chromatin, indicating its mixed role as a pioneer factor that opens chromatin, and a TF that binds already accessible DNA. This result is similar to those described for other pioneer factors, such as neurogenic factor Ascl1 (62) or pluripotency factors Oct4, Sox2, Klf4 (63). Similar to previous reports (14,64), we observed a large overlap between the MyoD-bound sites in fibro-MyoD and myoblasts (**Figure S6**). However, MyoD appears to be limited in its capacity to bind to all of its potential targets in fibro-MyoD cells, and thereby binds and opens only a fraction of myoblast-specific sites, consistent with previous studies suggesting that MyoD’s role as a pioneer factor is limited (12,19). This may partially explain the incomplete transdifferentiation in our system. Importantly, the MyoD binding signal in myoblasts at myoblast-specific chromatin sites that remained closed in fibro-MyoD cells, compared to the sites that did become accessible, MyoD was bound more weakly or not at all, even in myoblasts.

In addition to the reduced MyoD binding at non-reprogrammed chromatin sites in fibro-MyoD cells, we found that these sites displayed an enrichment of binding motifs for SAND-domain factors, a family that includes AIRE, GMEB1 and SKI (**Figure 3A**). Since SAND-domain factors were minimally expressed in fibro-MyoD cells, this finding points to an attractive possibility for improving reprogramming efficiency. AIRE is known to bind unmethylated H3K4 molecules, and it has been hypothesized to play a role in transcriptional regulation via chromatin remodeling, although the precise molecular mechanisms are not well understood (65). GMEB1 is known to interact with CBP (CREB Binding Protein), which can open closed chromatin sites (66). SKI can convert quail non-muscle cells into muscle cells (67,68). Therefore, we hypothesize that increasing expression of these SAND-domain proteins could help open a subset of myoblast-specific sites not reprogrammed by MyoD alone. Our quantitative approach for analyzing chromatin changes during transdifferentiation, based on classifying genomic regions with different chromatin reprogramming levels (CRLs) also suggests that increasing the nuclear concentration of MyoD or its co-factors could lead to higher CRL scores for myoblast-specific DHS sites. This is consistent with the finding that increasing nuclear localization of MyoD improves reprogramming efficiency (69). However, increasing MyoD to levels that are not physiologically normal could also lead to off-target binding events and improper reprogramming, as discussed below.

Chromatin-remodeling factors are another class of proteins that may improve cellular reprogramming. Our classification analyses indicate that the pre-existing repressive mark H3K27me3 is moderately predictive of chromatin opening at myoblast-specific DHS sites, while pre-existing active marks (H3K4me1, H3K4me2) are similarly enriched at both reprogrammed and non-reprogrammed sites (**Figure S10**). This suggests that inducing a “poised” chromatin state (59) at myoblast-specific DHS sites prior to MyoD overexpression, by targeted deposition of repressive mark H3K27me3, could lead to increased MyoD binding and improved reprogramming efficiency. Our observations are in agreement with the pre-existing enrichment of H3K27me3 at sites bound by NeuroD1 during induced neuronal differentiation (57), and by pioneer factor PU.1 during HDACi-induced remodeling (58). Similar to H3K27me3, another repressive mark, H3K9me3, was also found to be enriched along with active marks H3K27ac and H3K4me1, at sites bound by pioneer factor Ascl1 during transdifferentiation of mouse embryonic fibroblasts to neuronal cells (62). Interestingly, during initial stages of pluripotency reprogramming of fibroblasts, large H3K9me3-marked chromatin domains were reported to prevent binding of pioneer factors Oct4, Sox2, and Klf4, diminishing overall conversion efficiency (63). Our study did not find any discriminatory capacity of H3K9me3.

We also observed enrichment of active chromatin marks at fibroblast-specific sites that do not close down in fibro-MyoD cells. If this relationship is causal, then erasing active marks in some non-reprogrammed fibroblast-specific DHS sites could facilitate closing down of regulatory sites to shut off fibroblast-specific genes. Our findings highlight the need for genome-wide reprogramming studies to explore the concerted effects of other TF master regulators and chromatin remodelers in transdifferentiation. One such study observed that expression of BAF60C, a component of the SWI/SNF chromatin remodeling complex, induces myogenic priming in MyoD-mediated reprogramming of human ES cells (70). We note that the enrichment of active chromatin marks at sites that fail to close down during transdifferentiation could also be due to the fact that fibroblast-specific TFs, which might target these regions, are encoded by some of the genes not appropriately repressed. While possible, this hypothesis is not supported by our classification analyses (**Figure 3B, Figure S9B**).

Our DHS pattern analyses also revealed chromatin opening at off-target sites, which resulted in a total of 6,792 fibro-MyoD-specific ‘010’ DHS sites. Only about 6% of these sites show MyoD binding in fibro-MyoD, suggesting that off-target chromatin remodeling is mostly indirect. These off-target DHS sites are scattered randomly throughout the genome. In addition, they do not appear to have a significant effect on gene expression, as we only found seven genes that are ‘improperly reprogrammed’ in the sense that they are significantly upregulated in fibro-MyoD compared to both fibroblast and myoblast cells. These misprogrammed DHS sites might be attributable to supraphysiological levels of MyoD.

In summary, our study revealed a continuum of chromatin remodeling changes genome-wide and showed that chromatin remodeling deficiencies are associated with global transcriptional reprogramming bottlenecks during MyoD-induced transdifferentiation. We identified potential explanations for the incomplete reprogramming at the chromatin level, and suggest mechanisms to improve the process. Our approach for genome-wide analysis of the efficiency of cellular transdifferentiation driven by a TF master regulator can be applied to any transdifferentiation system, and will likely be particularly useful for characterizing transdifferentiation systems with relatively low epigenetic conversion efficiency.

### AVAILABILITY

The PBM data on human and murine TFs used in this study are available from the UniPROBE database (http://the_brain.bwh.harvard.edu/uniprobe) and the DREAM5 challenge (http://hugheslab.ccbr.utoronto.ca/supplementary-data/DREAM5/. GEO accession code: GSE42864). The histone modification, CTCF and Ezh2 ChIP-seq data corresponding to the normal human dermal fibroblast (NHDF) cell line were downloaded from the ENCODE repository and are available in the NCBI GEO database with the following accession codes: GSM733662 (for H3K27ac), GSM733745 (for H3K27me3), GSM733733 (for H3K36me3), GSM1003526 (for H3K4me1), GSM733753 (for H3K4me2), GSM733650 (for H3K4me3), GSM1003554 (for H3K79me2), GSM733709 (for H3K9ac), GSM1003553 (for H3K9me3), GSM1003486 (for H3K20me1), GSM1003505 (for H2A.Z), GSM733744 (for CTCF), and GSM1003550 (for Ezh2). The MyoD ChIP-seq data on primary myoblasts was obtained from (33) and is available in GEO (accession code GSE50415). The fibro-MyoD ChIP-seq data, as well as the RNA-seq and DNase-seq data used in this study, can be accessed in GEO database using the accession code GSE93268. This GEO SuperSeries includes GSE93257 (DNase-seq data), GSE93263 (RNA-seq data), and GSE93258 (fibro-MyoD ChIP-seq data). The DNase-seq data for myoblasts were obtained from (20,22) and processed as described in Methods. The RNA-seq data for myoblasts were obtained from (71) and processed as described in Methods.

## ACKNOWLEDGEMENTS

We thank Andrew Allen, Sayan Mukherjee and Barbara Engelhardt for helpful discussions, and the Duke Sequencing Core for library preparation and sequencing assistance.

## AUTHOR CONTRIBUTIONS

R.G., G.E.C. and C.G. conceived and supervised the study. R.G., G.E.C and D.M. conceived the computational analyses. D.M. performed the computational analyses. L.S. performed the high throughput sequencing experiments. L.S. and L.E. performed data analysis. A.K. and J.K. performed the transdifferentiation, immunostaining and cytometry experiments. K.T. and M.E. grew and provided primary human myoblast culture. R.G., G.C., and D.M wrote the manuscript, with assistance from all authors.

